# Humans use a local code for tactile perception

**DOI:** 10.1101/2019.12.12.874206

**Authors:** Arindam Bhattacharjee, Christoph Braun, Cornelius Schwarz

## Abstract

Humans are classically thought to use either spectral decomposition or averaging to identify vibrotactile signals. These are general purpose ‘global’ codes that require integration of the signal over long stretches of time. Natural vibrotactile signals, however, likely contain short signature events that can be detected and used for inference of textures, instantaneously, with minimal integration, suggesting a hitherto ignored ‘local code’. Here, by employing pulsatile stimuli and a change detection psychophysical task, we studied whether humans make use of local cues. We compared three local cues based on instantaneous skin position and its derivatives, as well as six global cues, calculated as summed powers (with exponents 1,2, and 3) of velocity and acceleration. Deliberate manipulation of pulse width and amplitude (local+global) as well as pulse frequency (global) allowed us to disentangle local from global codes. The results singled out maximum velocity, an instantaneous code, as a likely and dominant coding variable that humans rely on to perform the task. Comparing stimuli containing versus lacking local cues, demonstrated that performances exclusively using global cues are rather poor compared to situations where local ones are available as well. Our results are in line with the notion that humans not only do use local cues but that local cues may even play a dominant role in perception. Our results parallel previous results in rodents, pointing to the possibility that quite similar coding strategies evolved in whisker and finger tactile systems.

**Significance statement:** The brain is believed to select coding symbols in sensory signals that would most efficiently convey functionally relevant information about the world. For instance, the visual system is widely believed to use spatially local features, like edge orientation, to delineate a visual scene. For the tactile system only global, general purpose coding schemes have been discussed so far. Based on the insight that moving contacts, characteristic for active touch, feature short-lived stick-slip events, frictional movements that transfer fair amounts of texture information, one should expect the brain to use a temporally local code, extracting and instantaneously analyzing short snippets of skin movement. Here, we provide the first analytical psychophysical evidence in humans that this indeed is the case.

## Introduction

A sensory signal can transmit information about the world using either local or global variables. A classical dispute in vision research was between one view that early vision essentially is a set of global linear filters (Campbell and Maffei, 1974) and an opposing view that interpreted neurons as local feature detectors (Barlow, 1972; Hubel and Wiesel, 1968). In research on the tactile sense, local coding was rarely considered – at least for pure temporal coding of texture (roughness). Global coding schemes like spectral decomposition and finding the ‘best frequency’, as well as signal averaging to come up with ‘intensity’, dominated the thinking in tactile research of finger/hand related perception (LaMotte and Mountcastle, 1975; Luna et al., 2005; Yoshioka et al., 2001). However, insight from tribology and recent research in whisker-based sensation and perception raised the possibility that vibrotactile signals contain local features (i.e. short-lasting events) that can be extracted and these features contain large amounts of texture information (Schwarz, 2016) (Fig. 1A). In the whisker system, evidence supporting local codes come firstly from biomechanical studies describing prominent stick-slip movements (Oladazimi et al., 2018; Ritt et al., 2008; Wolfe et al., 2008), secondly from neuronal coding on the ascending tactile pathway, revealing that neuronal spikes respond to local features in the tactile stimulus (Chagas et al., 2013; Jones et al., 2004; Maravall et al., 2007; Petersen et al., 2008), and finally from perceptual studies showing that pulsatile whisker deflections, devoid of local cues, are poorly discriminated (Gerdjikov et al., 2018, 2010; Waiblinger et al., 2015a, 2015b). Evidence pointing to this direction are also available in the fingertip system. Papillary ridges have been shown to undergo complex shear during lateral movement (Delhaye et al., 2016), the sensory consequences of which may reach perception (Barrea et al., 2018). Further, characteristics of vibrotactile signals as well as skin deformation when tapping support the possibility that tactile signals are processed and perceived instantaneously (Johansson and Birznieks, 2004; Lawrence et al., 2000; Pruszynski and Johansson, 2014; Weber et al., 2013).

**Fig. 1.**
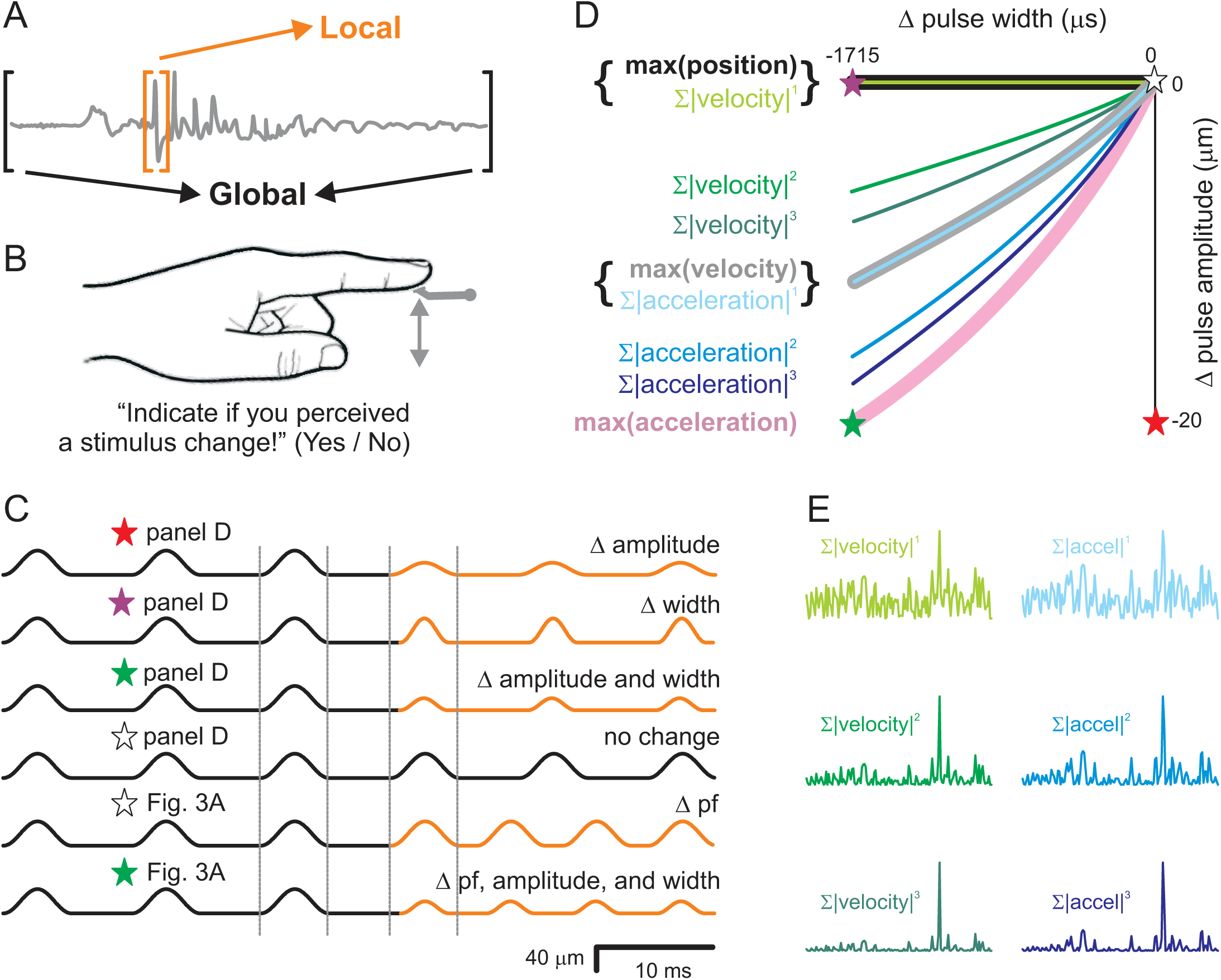
Definition of local coding and methods. **A**. The difference of local vs. global in the time domain of a vibrotactile signal. Local coding is defined as focusing on short-lived events in the vibrotactile signal, which can be extracted and analyzed (near-) instantly (golden bracket). Global coding on the other hand is an integrative mechanism, analyzing a long stretch of the vibrotactile signal using averaging or frequency decomposition (black bracket). **B**. In this study skin indentation of an actuated disk of 2.9 mm diameter at the fingertip of the left index finger was used. **C**. Participants received pulsatile stimuli consisting of single-period sinusoids separated by zero movement. They were instructed to indicate detection an abrupt change (orange) in either pulse waveform (Δ *amplitude*, Δ *width*; experiment 1, upper four stimuli), pulse frequency (Δ*pf*, experiment 3, 5^th^ stimulus), or both (experiment 2, 6^th^ stimulus). The stimulus containing no change (4^th^ stimulus, all black) was presented in all experiments in 50 % of the trials. All traces are to scale and demonstrate the extremes of the stimulus space used (see stars in panel D and Fig.3A). The changes shown are: amplitude: from 40 to 20 µm; width: from 5882 to 4167 µs; pulse frequency: from 90 Hz to 135 Hz. Note, for demonstration purposes only a few pulses around the change are shown. **D**. Definition of coding variables and stimulus space. Stimulus space spanning the change of pulse waveforms presented in experiment 1 is shown. Abscissa: change of pulse width, Ordinate: change of pulse amplitude. The reference stimulus (‘no change’) is at (0|0). Stimuli located on the nine iso-lines shown do not change the coding symbol indicated on the left (color of line and coding symbol matches). Iso-lines of intensity variables (global) are indicated by thin lines. Shades of green indicate velocity-based, while shades of blue indicate acceleration-based variables. Instantaneous kinematic variables (local) are plotted using thick lines. Note that two pairs of iso-lines (curly brackets) are congruent. Asterisks on the corners mark the extreme stimulus changes demonstrated in C. **E**. Intensity variables integrate long signal stretches. However, non-linear elements in their definition as well as differntiation, can change the weights with which local features enter the value of a global variable. An arbitrary velocity signal and respective first derivative (accel=acceleration) is shown in the upper row and taken to different powers (1,2, and 3; along columns). Higher derivative / higher power intensity formulations are increasingly dominated by local features in the signal.

Here we used psychophysics based on pulsatile fingertip skin indentation to demonstrate for the first time, that humans use local codes. The experimental idea was that the change of single pulses’ waveform without changing pulse frequency is a manipulation of local and global variables, whereas manipulation of pulse frequency without waveform changes is a manipulation of global variables (Waiblinger et al., 2015a) alone. Our results strongly indicate that humans do use a local code and may even use it as the dominant code in addition to the classic global ones.

## Results

We established a Yes/No change detection task. In each trial two 500 ms long pulse trains, concatenated in a seamless way, were presented as vertical indentations to the left index fingertip of participants (Fig. 1BC). After experiencing a single presentation of such stimuli, participants indicated whether they perceived (Yes/No) a stimulus change in the middle of the stimulus presentation (Fig. 1C; black stimulus [4^th^ from top]: correct response – No; all other exemplary stimuli: correct response - Yes).

### Experiment1

Our first aim was to elucidate systematic performance deficits with deliberate manipulation of pulse width and amplitude (as shown in the four upper stimuli of Fig. 1C). In this experiment the pulse frequency of all the trials was kept constant at 90 Hz, and thus, did not provide any cue of stimulus change. In principle, the local changes to pulse waveforms can bring about changes in the global intensity variable as well. The amount of these possible changes, however, depends very much on how intensity is defined. In the literature a variety of intensity definitions have been used, such as linear integration (‘mean speed’), or non-linear integration (‘power’) of different kinematic derivatives (position, velocity, acceleration). In fact, an accepted standard measure of intensity does not exist. Therefore, instead of using one fixed intensity definition, we decided to use an array of them. Since in principle there are infinite ways of defining intensity – we opted for an array of definitions that firstly vary the code’s characteristics in a systematic way across the stimulus space; presumably, one of the coding definitions would capture specific performance deficits in stimulus space related to the sought-after unknown coding variable. Secondly, we incorporated a non-linear element (taking the signal to the power of >1 before summation) in the definition of intensity that systematically varies the emphasis of local features within the vibrotactile signal. This was done to address the possibility that local coding could be realized by a mathematical integration with non-linear preprocessing as well (as opposed to the truly local, instantaneous feature extraction that we portrayed above).

Intensity was thus defined as the sum (or equivalently as the mean) of the first and second stimulus derivative (i.e. velocity and acceleration) taken to the powers 1, 2, or 3. This choice satisfied the spread in stimulus space. Keeping each of those intensities constant (under the assumption that only pulse amplitude and width are varied), yielded an array of so-called ‘iso-lines’ fanning out from the origin (no change) into relevant sections of the space spanning possible stimulus changes (tagged ‘stimulus space’ for short throughout this report; Fig 1D). To illustrate the stimulus space, the extreme stimulus changes (in the corners of the stimulus space) are marked with a star and the respective waveforms can be looked up in panel C. Thick lines represent iso-lines that keep one of the local variables constant (maximal pulse position, velocity, and acceleration, colored black, gray and pink, respectively) while thin lines represent iso-lines of global variables (shades of green: velocity-based; shades of blue: acceleration-based). It is important to note that two of the local iso-lines overlapped with global ones (‘maximum position’ with ‘mean speed’, and ‘maximum velocity’ with ‘mean absolute acceleration’). (These overlaps required extra experiments to disentangle them – cf. experiment 2 below).

Moreover, our array of (global) intensities was designed to cover as well different degrees of emphasis on local features. To visualize what that means we demonstrate an arbitrary signal containing a shallow local feature in the upper left corner of figure 1E. The signal is differentiated along rows, and is taken to increasing powers along columns. From this demonstration one can easily appreciate that low derivative / low power signals (e.g. *speed* = |*velocity*|^1^) de-emphasize local features, whereas high derivative / high power signals increasingly emphasize them (e.g. *absolute cubic acceleration* = |*acceleration*|^3^).

Armed with this analytic stimulus design, we set out to measure psychophysical performance of 10 participants on stimulus changes located on the iso-lines depicted in figure 1D. The reference stimulus is located at the origin of the coordinate system spanning changes in pulse amplitude and width (pulses of 40 µm amplitude and 5882 µs width). The comparison stimulus (seamlessly concatenated to the reference stimulus) was picked in each session from a different iso-line (15 equidistant pulse widths between 4167 (easy) and 5882 µm (difficult)). Each session comprised 840 trials with 50% no change stimuli (comparison = reference = 5882 µm) and 50% change stimuli (comparison ≠ reference, i.e. 30 trials per 14 pulse widths). We refrained from testing global variables based on stimulus ‘position’ because the iso-lines for position-based intensities are located in a part of the stimulus space where human ability of detection of change was superb whenever a stimulus differed ever so slightly from the reference (in the second quadrant, where amplitude increased and width decreased, Fig. 1D). This stimulus area, thus, provided little experimental leverage to gain insights into the tactile codes used (the respective preliminary experiments are not shown).

The typical performance of a single participant is shown in figure 2A. It can be seen that the participant shows a remarkably well isolated deficiency centered on the iso-line of maximum velocity (gray). This observation generally held true in all ten participants. Comparing effect sizes of the thresholds obtained from all ten participants on the maximum velocity iso-line compared to all other iso-lines showed that the poorest performance was located around the iso-line of maximum velocity (AUC comparing to 7 iso-lines in the order listed in Fig. 1D: [0.98, 0.82, 0.53, 0.5, 0.83, 0.91, 0.90]; corresponding p values in t-tests were [0.015, 0.044, 0.624, ∼, 0.357, 0.049, 0.055]; n=[10,9,9,∼,9, 10, 10]). The best fit logistic functions to the pooled data from ten participants and the respective thresholds are shown in Figure 2BC. It is worth mentioning that in general the psychophysical test was subjectively described as ‘demanding’ by all participants. In fact one person failed on the task (producing no psychometric fit passing p=0.5 in any of the sessions) and was excluded from all the analyses. Two other persons dropped out after a few sessions without giving reasons for their decision. Reflecting the challenging nature of the task we observed a consistently high false alarm rate of [0.12, 0.18, 0.16, 0.14, 0.09, 0.04, 0.07] on the 7 iso-lines as listed in figure 1D.

**Fig. 2.**
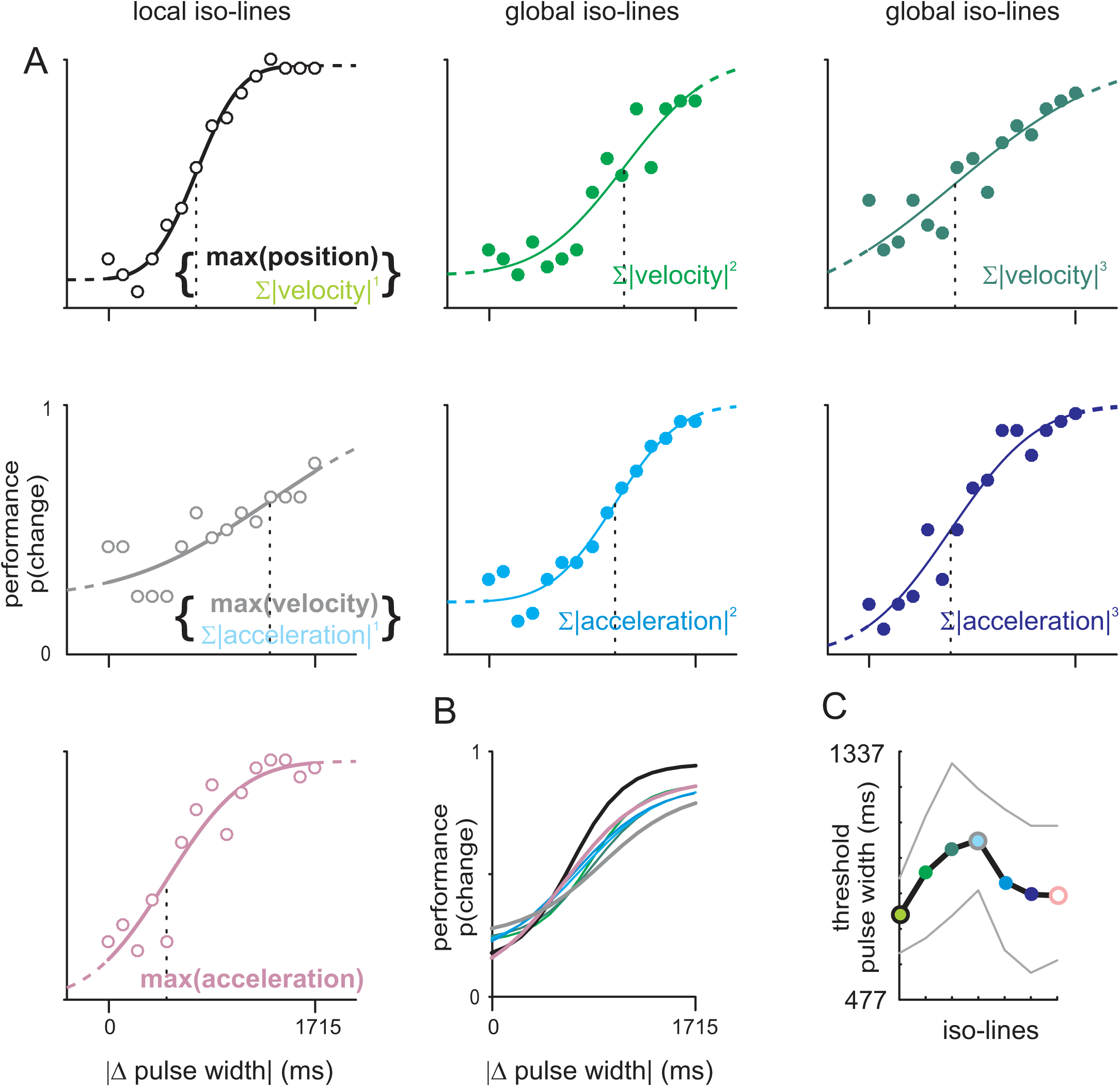
Experiment 1. Detection of pulse waveform changes. **A**. Data and logistic fits to the performance of one participant on all iso-lines shown in Fig. 1D. Same axes scaling in all sub-panels. **B**. Logistic fits using pooled data of all 10 participants. **C**. Thresholds and 95% confidence intervals expressed as pulse width for all isolines. Performance on ‘maximum velocity’ iso-line is poorest across all participants. Colors match throughout all panels. Thick lines / hollow symbols: iso-lines of local variables; Thin lines / filled symbols: iso-lines of local variables. Statistics see text.

### Experiment 2

So far an unambiguous conclusion about the role of local vs. global variables in mediating the poor performance close to the maximum velocity iso-line cannot be reached because the ‘maximum velocity’ iso-line was congruent with that of the global variable ‘mean absolute acceleration’. To disentangle local from global variables, here we implemented the procedure of experiment 1 with an additional manipulation of pulse frequency (*pf*; cf. the bottom two stimuli in Fig. 1C). Pure changes of pulse frequency (4^th^ stimulus from the top in Fig. 1C) do not affect instantaneous variables, because the pulse waveform is identical. They do however change the iso-lines of all global variables, which shift downward toward larger decrements in pulse amplitude to balance out the presence of additional pulses. The shift introduced by a change in pulse frequency Δ*pf* = 15 *Hz* is demonstrated by the move from the blue [Δ*pf* = 0] to red [Δ*pf* = 15*Hz*] curves in Fig. 3A. The thick gray line is the ‘maximum velocity’ iso-line, which does not move when the Δ*pf* cue is added.

**Fig. 3.**
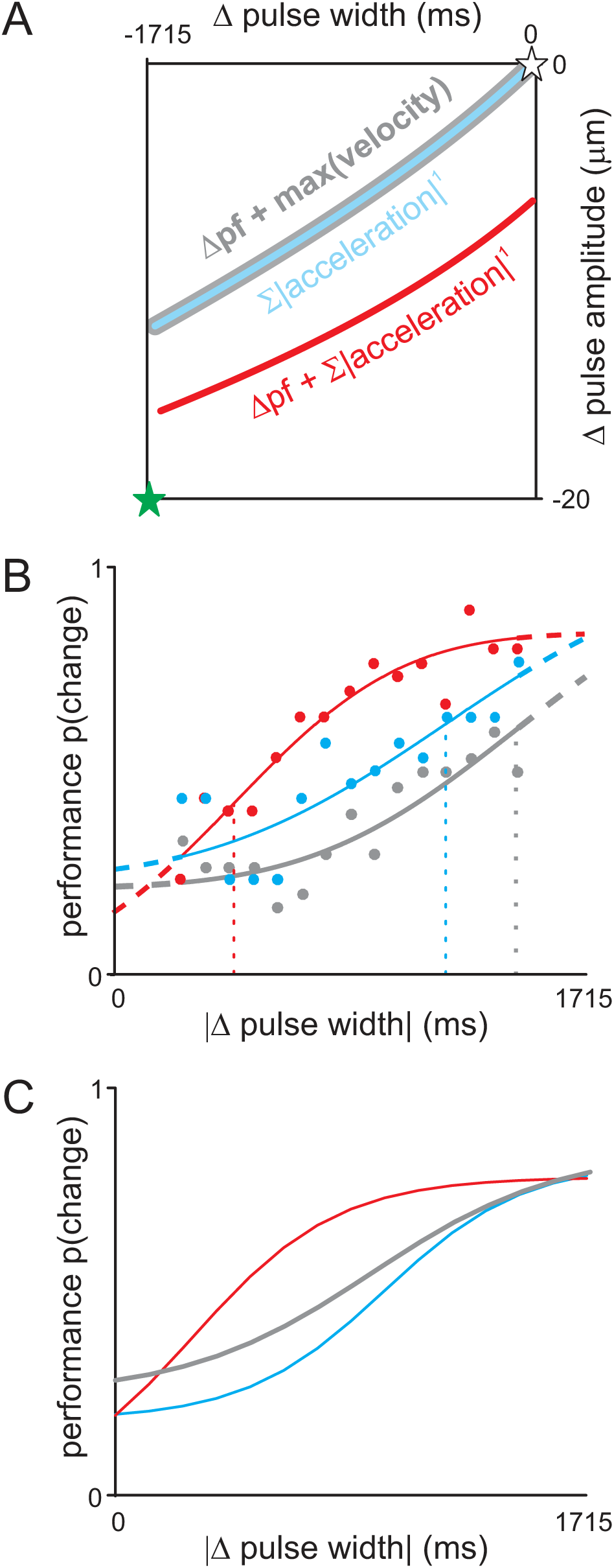
Experiment 2. Disentangling the congruent iso-lines related to ‘maximum velocity’ and ‘mean absolute acceleration’. **A**. Iso-lines with (gray and red) and without (blue) changes in pulse frequency (Δ*pf*). Iso-lines of local variables do not change with adding pulse frequency cues while those of global variables do. The gray curve is the unchanging iso-line of maximum velocity with and without Δ*pf*. The blue and red curves are the iso-lines of mean absolute acceleration with and without Δ*pf*, respectively. Effectively then, the addition of Δ*pf* disentangles the two isolines. For examples of stimulus waveforms refer to figure 1C. **B**. Data and logistic fits for the same participant shown in figure 2B. **C**. Logistic fits for the population of 9 participants. Poor performance is pegged to maximum velocity. There is a slight non-significant improvement in performance when adding Δ*pf* (blue to gray). However, shifting the mean absolute acceleration iso-line with Δ*pf* improves performance significantly (blue to red). Colors in all panels match. Thick lines: iso-lines and respective performance of local variables; thin lines: same for global variables

Pulse frequency is a cue in itself - independent from the abovementioned instantaneous kinematics (local) and intensity (global). In fact, as mentioned above, frequency is the second candidate variable, next to intensity, for global coding. Therefore, pulse frequency would be expected to change the detection rate by itself; however, this effect should be the same for all iso-lines. Our choice of Δ*pf* = 15 *Hz* (reference value 90 Hz, comparison value 105 Hz) balances a low perceptual effect (mean increment of the probability to report a ‘change’ 0.15, SD 0.08; as measured in experiment 3) with a large-enough shift of the iso-line to capture eventual differences in performance. Figure 3B demonstrates the performance of the same participant as shown in figure 2A on stimuli from the three iso-lines depicted in Fig. 3A. The effect of pure global stimuli (frequency and intensity) is reflected in the difference between the blue and gray line. It shows a non-significant mean change in the threshold from 1101 µs (SD 194 µs) (Δ*pf* = 0) to 1272 µs (SD 270 µs) (t=-2.084, n=8, p=0.08; AUC=0.13). However, the performance on the shifted iso-line of ‘Δ*pf* + mean absolute acceleration’ (red) is significantly improved to a mean change of threshold of 764 µs (SD=368 µs, t=2.868, n=8, p=0.024; AUC=0.99). Figure 3C shows the logistic fits for the pooled performance from all 9 participants. In summary, we found that poor performance was pegged to the local ‘maximum velocity’ iso-line, and did not move together with the global ‘mean absolute acceleration’ iso-line.

### Experiment 3

The participants in experiment 2 were in addition tested on pulse trains that exclusively changed in Δ*pf* (cf. the gray stimulus in the schematic of Fig. 1C; only data from 8 of 9 persons are reported here as from the 9^th^ we did not get any logistic fit that crossed a probability of correct responses of 0.5). Figure 4A presents the logistic fit to the pooled trials of this population. The success of 8 out of 9 participants to detect pulse frequency changes demonstrates that humans in principle are able to use global variables as a basis for their decision. The drop out of one participant and the subjective difficulties observed in others, however, raised the question, how well the successful participants were able to use them, and how their performance compared to situations in which they had additional access to local cues. To this end we scaled the performance on experiment 3 (Δ*pf* cue and respective intensity cues, thick red lines in figure 4B) and those obtained in experiment 1 (intensity cues and local cues, colored lines in figure 4B) to each of the 6 variants of intensity (one variable per sub-panel). In the vast majority of these direct comparisons the performance measured in experiment 1 was shifted to the left and showed far lower thresholds of respective intensive variables as compared to the Δ*pf* stimuli used in experiment 3. That is, generally, the performance when only global cues were present was rather poor compared to the performance with access to the same intensity variable and additional local ones. While this held true for all almost all intensity variables, it was also clear that the definitions using higher derivative and powers seem to fare better in this comparison. In the plot of ‘mean cubic absolute acceleration’ (dark blue), two of the curves obtained in experiment 1 showed even slightly higher thresholds than the ones obtained in experiment 3. This is summarized in figure 4C, which plots the relative thresholds obtained in experiment 3 (red stars) and experiment 1 (dots colored with respect to the global variable, same colors as in panel B) on one and the same scale (note that abscissa scales in the sub-panels of Fig. 4B differ as they are individually normalized to the highest value obtained in experiment 3). In summary, relative thresholds yielded by different intensity variables are all similar, a fact that shows that none of these variables stands out in its importance for perception. Moreover, the perceptual effects of intensity variables, particularly the definitions based on lower derivatives and power, are far inferior compared to local cues, as experiment 3 (changes in intensity and pulse frequency) typically resulted in far higher thresholds than those obtained in experiment 1 (changes in intensity and instantaneous kinematics). This was so, despite the fact that participants might have additionally exploited pulse frequency cues from stimuli in experiment 3, which were not present in experiment 1. Finally, it can be appreciated that intensity definitions that are based on the higher kinematic derivative (i.e. acceleration) and higher power (i.e. power of 3) achieved thresholds that are on par with local variables. This chimes well with the fact that these higher order definitions of intensity are increasingly dominated by local features (demonstrated in Fig. 1E). We interpret this match as indicating that our array of intensity definitions exhausts the range of possible formulations that truly represent global features.

**Fig. 4.**
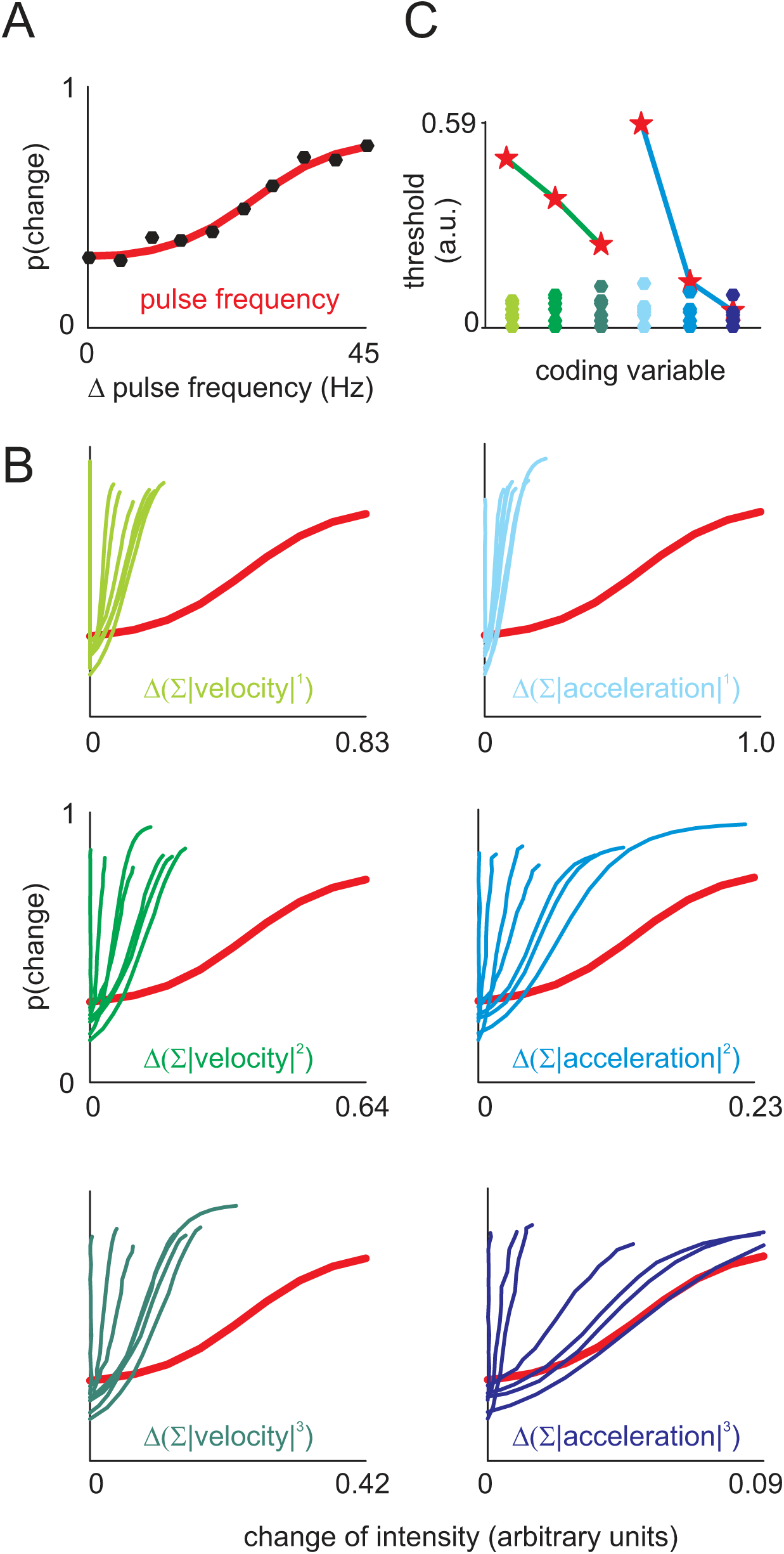
Performance with and without local cues. **A**. Performance on stimuli using pure global cues. i.e pure pulse frequency changes (Δ*pf*, experiment 3; cf. 5^th^ stimulus in Fig. 1C). **B**. Replotting data from A (red lines) and data obtained from the performance on seven iso-lines in experiment 1. Each sub-panel plots the data from 7 iso-lines, scaled according to one of six intensity variables. Each intensity variable in the six sub-panels is normalized and scaled to the maximum of that variable found in the Δ*pf* stimuli. **C**. Thresholds of all logistic fits to the data shown in B, now all normalized to the same scale. Red stars relate to red lines in B (experiment 3, Δ*pf* stimuli), colored dots relate to the lines in B of the same color (experiment 1, pulse waveform changes). Note the descending threshold obtained when basing the performance on increasing power (red stars, green line: velocity-based intensity, blue line: acceleration-based intensity). This comparison across scales suggests that intensity definitions using higher powers (weighing local features higher) increasingly represent performance as true local variables do.

## Discussion

In this study, we present novel psychophysical evidence that humans use local instantaneous codes in addition to the classical integrative global codes for tactile discrimination. Using manipulations of pulsatile skin-indentation stimuli, we found that taking ‘maximum velocity’ out of the pool of available cues led to poor performance on a Yes/No psychophysical change detection task. Finally, performance on pulse frequency changes (engaging exclusively global coding mechanisms), was found to be significantly inferior when comparing them to performance on sets of stimuli that contained the same changes in global parameters but in addition allowed access to local ones. This last finding speaks in favor of the notion that humans not only use local codes, but that they may even rely on them as the dominant source of information feeding tactile perceptual decisions.

### Is it appropriate to use pulsatile stimuli to disentangle local from global coding?

Classically, sinewave stimuli have been used to disentangle ‘intensity’ (sine amplitude) from ‘frequency’ (sine frequency) (LaMotte and Mountcastle, 1975). This approach, however, only allowed addressing global codes because changes in sine amplitude and/or frequency would always concomitantly change the instantaneous signal value, i.e. local coding variables. Thus, for the quest to find out whether local vs global codes are used, the usage of simple sinusoidal stimuli is prohibitive. In order to investigate local features, stimuli need to be manipulated locally in the time domain (Waiblinger et al., 2015a, 2015b). Pulsatile stimuli are an attractive tool for this purpose as they are midway between sinusoids and broad band stimuli featuring spectra that contain a perceptually important peak of power at the base frequency (i.e. the pulse frequency) and allow for simple and systematic changes of individual (local) pulses. Moreover, by introducing, eliminating, or shifting pulses around in time one can easily introduce changes in global cues (frequency, intensity) while keeping local ones (pulse waveform) identical (Gerdjikov et al., 2018). However, there are two major problems with this strategy. Firstly, the success in isolating local changes hinges on the definition of the global code. Certain changes of pulse waveforms may change one global variable but not another. Part of the problem is that in the literature, intensity codes have never been standardized – for instance many studies used signal power (squared sum, e.g. Hipp et al., 2006), others used a simple sum of the absolute signal (e.g. Arabzadeh et al., 2003), etc. We addressed this problem by testing not only one possible intensity code but defining a whole array of them. The iso-lines of our selection of intensity variables cover a large part of the stimulus space (Fig. 1D). Additionally, our strategy to calculate intensity based on different powers of the signal helps to formulate global codes that do or do not emphasize local features (low vs. high derivatives/powers) (Fig. 1E). In fact, our intensity variable that most strongly emphasizes local features, ‘mean cubic absolute acceleration’, yields performance estimates that begin to match the ones obtained with true (instantaneous) local codes (Fig. 4BC), indicating that our sample of intensity codes entirely covers the space spanned between (true) local coding, global coding with emphasis on local features, and (true) global coding without that emphasis. This seems important because (in principle) infinite other definitions of intensity are possible, which, however, are likely to be close to one of our present definitions in terms of iso-line location and bias toward reporting local features.

The second issue with pulsatile stimuli is their spectral composition. Any modification of pulse waveform (as done in experiment 1) will keep the signal’s base frequency but will introduce wide ranging spectral changes at higher harmonics. It follows that what we call ‘local coding’ in the time domain relates to differences of spectral patterns at higher harmonics, and therefore is ‘global’ in the frequency domain. From these consideration two aspects are noteworthy: First, our terms ‘local’ and ‘global’ strictly pertain to the time domain, where a pulse is local and a single frequency (like the base frequency) is global. Second, our results cannot be easily compared to studies that use definitions of ‘frequency coding’ based on more than single spectral elements(e.g. Manfredi et al., 2014), because frequency coding defined that way may in fact involve a fair amount of local coding in the time domain.

There are several hints that in primates and rodents the base frequency of a pulsatile code plays a minor role for tactile perception. Rats show exactly the same performance in detecting changes of pulsatile stimuli irrespective of whether intensity or pulse frequency is changed (Waiblinger et al., 2015a). In rat primary somatosensory cortex spike counts relate better to tactile discriminations than timing of spikes (Gerdjikov et al., 2018). Monkeys perform frequency discrimination on pulsatile stimuli independent of the rhythmicity of pulses (Salinas et al., 2000), and humans perceive frequency based on the longest gap between bursts of pulses irrespective of the absolute number of pulses (at least in lower frequency ranges; Birznieks and Vickery, 2017). Our experiments add to the view that spectral analysis may play a rather inferior role in tactile perception. Our quantitative comparison of performance obtained with pure pulse frequency changes in comparison to stimuli containing local cues (Fig. 4), demonstrated the inferiority of base frequency cues to determine perception.

### What is the functional advantage of a local code?

The search for local coding is motivated by the assumed presence of frictional movements in objects that engage in moving contact (Schwarz, 2016). Moving contact is at the heart of tactile processing, as palpation movements are indispensable for the perception of fine textures (Hollins et al., 2001; Hollins and Risner, 2000; Skedung et al., 2013). Prototypical expressions of frictional contact are stick-slip events - short lived elastic deformations of the contacting materials coming about by sticking to surface elements, storing energy in elastic deformation, and releasing them quickly into sudden slips when frictional force is overcome by the driving movement of the contact (Schwarz, 2016). In the rodent vibrissa-based tactile system, frictional slips have been shown to exist (Arabzadeh et al., 2005; Ritt et al., 2008; Wolfe et al., 2008), and to encode texture information (Oladazimi et al., 2018; Wolfe et al., 2008). Primary sensory cortex in these animals has been shown to be selective for slip-based tactile inputs (Jadhav et al., 2009). Stick-slip events are short-lived and therefore local in character, meaning that any tactile information they store is immediately available. An efficient decoding system making use of stick-slip events is supposed to have evolved under time-accuracy constraints, and thus, must be assumed to be local itself, i.e. make use of tactile information immediately after the reception of slip-based tactile signals. In support of this notion, manipulations of pulsatile whisker deflections in a psychophysical study, have provided strong evidence in favor of local coding in rats’ tactile system (Waiblinger et al., 2015a). In the fingertip system evidence about frictional movements is more scant, but by no means negligible: Papillary ridges undergo complex shearing deformations at the onset of a finger movement (Delhaye et al., 2016), and friction has been shown to be a determinant of roughness estimation – most of all for the discrimination of microscopic surface elements (Verrillo et al., 1999). Research on prehension has unearthed unequivocal evidence that sudden slip movement does occur in the skin, and is readily detected by humans to adjust grip forces (Johansson and Westling, 1987). Finally, so-called rate hardness (an instantaneous measure of change of force and speed when tapping surfaces) relates tightly to hardness perception suggesting that it may be based on local elements of the tactile signal as well (Han and Choi, 2010; Lawrence et al., 2000). On the anatomical level, the presence of papillary ridges in glabrous skin, beset by saliently structured rows of ridge-associated Meissner corpuscles and Merkel cells (Cauna, 1954) has rarely been attempted to be incorporated into a functional hypothesis (Gerling and Thomas, 2008). Biomechanical generation of stick-slip in ridges, and specific reception of these by the mentioned rows of ridge-associated receptors is a hypothesis that may help to unearth such functional links in the future. Currently, these are mostly indirect evidences speaking in favor of local coding, but due to lack of more detailed investigations it seems difficult to dismiss local coding as playing a critical role in fine texture perception in primates (Schwarz, 2016). Nevertheless, on the behavioral level, no evidence directly supporting this notion was available so far. Our present results are the first to provide systematic observations that can begin to fill this gap and demonstrate that kinematic features of vibrotactile signals represent a perceptually powerful coding object.

## Methods

### Participants

We recruited a total of 13 neurologically-healthy (self-reported) participants (age:20-35 years, median 27 years; 5 female). Two participants withdrew from the study without providing any reason. Our participant recruitment advertisement discouraged all individuals from participation if they were diagnosed with dyslexia (because it adversely affects tactile acuity, (Grant et al., 1999; Laasonen et al., 2001), diabetes (which could result in peripheral neuropathy and action potential conduction delays, (Hyllienmark et al., 1995), learning disabilities, nervous system disorders, or had any calluses or injury to the left index fingertip (the tested finger). Based on questions modified from the Edinburgh Handedness Inventory (Oldfield, 1971), we classified 12 participants as right handed. The study was approved by our institutional research ethics board; all the participants signed the informed consent form and were paid for their participation in the study.

To identify the stimulus feature that the participants used to perform the perceptual tasks, we required the participants to perform well so that we could generate their psychometric function for each task (see subsection perceptual task). If any participant performed poorly on any task (session) and failed to generate a viable psychometric function for those tasks (passing 50% correct), we ran them again on those specific tasks. Three participants completed the study in a single attempt, 4 participants redid 1 out of 7 tasks, and 3 participants redid 2 out of 7 tasks. One participant was disqualified from the study because total percent correct for all 7 sessions using stimuli from the 7 iso-lines in experiment 1 was less than 50%. The data of the disqualified participant and the participants who withdrew from the study are not included in the results section of this study.

### Vibrotactile stimulation

We applied passive vibrotactile stimuli (i.e. no finger movement) to the distal pad of the left index finger, using a plastic circular disc of 2.9 mm diameter attached to a galvomotor (Cambridge Technology, Massachusetts; model 6220H). The galvomotor was driven by a custom made amplifier, which reproduced highly precise displacements. We calibrated the displacements of the galvomotor using a laser distance estimator that is sensitive to displacements at the micron resolution. We used Matlab (Natick, USA) to generate the stimulus waveform and control the galvomotor movement by passing voltage waveforms through an analog output channel digitized at 40,000 samples per second at 12 bit resolution via a National Instruments PCI-MIO-16E-1 I/O board.

Participants’ arm rested on a platform which could be raised or lowered depending on the participants’ comfort. To prevent finger movements, the index finger was clamped in a finger housing using a cleft as a rest for the finger nail - in addition to a double-sided tape that affixed the plane of the fingernail to the ceiling of the housing. Once the testing finger was securely positioned, we used a tri-axis micromanipulator to adjust and attach the galvomotor to the distal pad of the left index finger such that the circular disc area was completely covered by the fingertip skin. During the experiment only the testing region of the fingertip touched the circular disc, we ensured that no other part of the galvomotor touched any part of the participants’ finger. We asked participants to trim the nail of their testing finger to prevent any possibility of their nail touching any part of the stimulator. The arm platform as well as the galvomotor platform were separate from each other and seated on an anti-slip and anti-vibration mat. The depth of indentation was defined as the position in which during very slow movement toward the skin the participants would first report touch-down. Using a micromanipulator the galvomotor was then proceeded to the null position at a depth of 1 mm. From there the stimulus pulses further indented the skin.

Pulsatile stimuli were constructed by using one period sinusoids (waveform of a sinusoid extracted from one of its minima to the next) as done before (Gerdjikov et al., 2010). Two manipulations were performed to change pulse width and amplitude. For pulse width, the sinusoids used for this procedure ranged in frequency between 170 and 240 Hz in steps of 5 Hz, resulting in 15 pulse waveforms that varied in pulse width between 1°s°/°170°Hz°=°4167°µs and 1°s°/°240°Hz°=°5882 µs. The pulse amplitude ranged between 20 and 40 µm. A third manipulation left pulse waveform untouched but changed pulse frequency, i.e. the inverse of interpulse intervals. Pulse frequency of the reference stimulus was always 90 Hz. It changed in 5 Hz steps to values up to 135 Hz. The stimulus was a seamless concatenation of two pulse trains both 500 ms in duration. The first, called reference stimulus, was a train at a pulse frequency of 90 Hz, a pulse amplitude of 40 µm, and a pulse width of 5.882 ms. The second train of pulses was either the same as the reference stimulus (i.e. ‘no change’) or one that was altered into one or several of the above-mentioned manipulations (i.e. ‘change’)- see Figure 1C (the traces are to scale but for purposes of visualization the traces contain only a few pulses pre- and post-change).

### Perceptual tasks and psychophysics procedure

The participants were instructed to indicate in a Yes/No fashion their decision about the absence or presence of the perceived stimulus change. This was done by pressing one of two buttons (Yes/No) on a wireless presenter clicker with their right hand. The intertrial interval started after the participant’s response and lasted 5 s. After each block of 280 trials (140 ‘no change’ trials), the participants took a minimum of 2 minutes break in which they were encouraged to stand up and walk around. Participants received feedback after each trial which was delivered through wireless headphones. In addition, after each 280 trial-block participants saw their performance as total percent correct for that block. During experiment 2 and 3, white noise was played out loud next to the tactile stimulator. Following the completion of experiment 2, when asked, none of the participants was aware that the pulsatile frequency (along with the pulse width and amplitude) of the target stimuli changed.

### Statistical analysis

To each participant’s performance (proportion reported “change”) in each task, using the dedicated analysis software *psignifit* (Wichmann and Hill, 2001), we fitted a mixture model cumulative normal psychometric function of the following form:

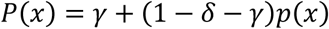

where p(x) is the sigmoid modelled as cumulative gaussian – which includes the threshold (i.e. the mean) and width (i.e. the SD) parameters, γ parameter is false alarm (lower asymptote), and δ parameter is the lapse rate (upper asymptote). The *psignifit* algorithm computed the maximum likelihood of each psychometric function generated from the combination of each of the above-mentioned parameters. By marginalizing over the γ, δ, and width parameters *psignifit* yielded a posterior probability density function (PDF) over the threshold parameter(θ), i.e. where the psychometric functioned crossed 50% correct. For each participant’s point estimate of their performance on each iso-line, we chose the stimulus value corresponding to the mode of the threshold PDF.

Next, to estimate the population mean for each iso-line, we implemented a Bayesian Hierarchical analysis (see Tong et al., 2016). We represented the participants’ threshold as normally distributed over the whole stimulus range (pulse width from 5882 down to 4167°µs) with unknown mean µ and standard deviation σ (ranging between 0.5 and 30). The probability of a participant’s data given (µ, σ) can be written as:

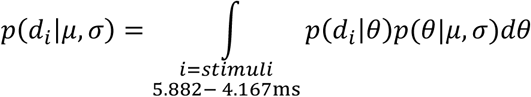

Here, the first term in the integrand is proportional to the threshold PDF of each participant (acquired from *psignifit*), because we assume a uniform prior over all possible values of θ (i.e. 5882 to 4167°µs). Thus, the likelihood of joint distribution of each (µ, σ) of complete dataset ‘*D*’ including all 10 participants can be represented as:

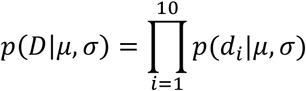

Finally, we marginalized over σ and obtained a PDF of the population mean; we report the mode of this distribution as the mean of the population along with the Bayesian 95% confidence interval.

### Experiment 1

This experiment exclusively used changes in pulse waveform and set the pulse frequency to 90 Hz throughout. We tested each participant on stimuli changes drawn from 7 iso-lines shown in Fig. 1D (seven sessions were performed, testing one iso-line per session). A session contained 420 ‘change’ and 420 ‘no change’ stimuli. The iso-lines contained stimuli from which we extracted 3 local (maximum position, velocity and acceleration) and 6 global (intensity) variables. The terms are given in Fig. 1D (the integration window was always the entire 500 ms stimulus, therefore we use sum and mean of the variable inter-changeably). In two instances an iso-line was found to be congruent with another (‘maximum position’ / ‘mean absolute velocity’ and ‘maximum velocity’ / ‘mean absolute acceleration’); hence, 7 iso-lines were tested. In one iso-line session the presentation was three blocks of 280 trials each, which contained all ‘change’ stimuli (140) and the same number of ‘no change’ stimuli trials in pseudorandom order.

### Experiment 2

The aim here was to disentangle performance on the ‘maximum velocity’ iso-line from that of the ‘mean absolute acceleration’ iso-line, using additional changes of pulse frequency (Fig. 3A). The change in pulse frequency was set to 15 Hz (i.e. from 90 to 105 Hz), a value that was far sub-threshold for all participants (see results). Otherwise the experimental conditions were as described for experiment 1.

### Experiment 3

Here stimulus changes were exclusively based on changes in pulse frequency (Δ*pf*). There were 9 ‘change’ stimuli and one ‘no change’ stimulus’, yielding 540 trials per session (30 trials per stimulus). Otherwise conditions were the same as for experiment 1.

## Acknowledgements

This research was supported by a grant from the Deutsche Forschungsgemeinschaft (SCHW577/14-1).

## Author contributions statement

AB designed and conducted experiments, analyzed data and wrote the paper. CB designed the experiments and wrote the paper. CS designed the experiment, analyzed data, and wrote the paper.

## Data availability statement

The datasets generated during and/or analyzed during the current study are available from the corresponding author on reasonable request.

## References

Arabzadeh E, Petersen RS, Diamond ME. 2003. Encoding of whisker vibration by rat barrel cortex neurons: implications for texture discrimination. J Neurosci 23:9146–9154.

Arabzadeh E, Zorzin E, Diamond ME. 2005. Neuronal encoding of texture in the whisker sensory pathway. PLoS Biol 3:e17.

Barlow HB. 1972. Single units and sensation: a neuron doctrine for perceptual psychology? Perception 1:371–394.

Barrea A, Delhaye BP, Lefèvre P, Thonnard JL. 2018. Perception of partial slips under tangential loading of the fingertip. Sci Rep 8:1–8. doi:10.1038/s41598-018-25226-w

Birznieks I, Vickery RM. 2017. Spike Timing Matters in Novel Neuronal Code Involved in Vibrotactile Frequency Perception. Curr Biol 27:1485-1490.e2. doi:10.1016/j.cub.2017.04.011

Campbell FW, Maffei L. 1974. Contrast and spatial frequency. Sci Am 231:106–115. doi:10.1038/scientificamerican1174-106

Cauna N. 1954. Nature and functions of the papillary ridges of the digital skin. Anat Rec 119:449–468. doi:10.1002/ar.1091190405

Chagas AM, Theis L, Sengupta B, Stüttgen MC, Bethge M, Schwarz C. 2013. Functional analysis of ultra high information rates conveyed by rat vibrissal primary afferents. Front Neural Circuits 7:190. doi:10.3389/fncir.2013.00190

Delhaye B, Barrea A, Edin BB, Lefèvre P, Thonnard J-L. 2016. Surface strain measurements of fingertip skin under shearing. J R Soc Interface 13:20150874. doi:10.1098/rsif.2015.0874

Gerdjikov T V, Bergner CG, Schwarz C. 2018. Global Tactile Coding in Rat Barrel Cortex in the Absence of Local Cues. Cereb Cortex 28:2015–2027. doi:doi: 10.1093/cercor/bhx108

Gerdjikov T V, Bergner CG, Stüttgen MC, Waiblinger C, Schwarz C. 2010. Discrimination of vibrotactile stimuli in the rat whisker system: behavior and neurometrics. Neuron 65:530–40.

Gerling GJ, Thomas GW. 2008. Fingerprint lines may not directly affect SA-I mechanoreceptor response. Somatosens Mot Res 25:61–76. doi:10.1080/08990220701838996

Grant AC, Zangaladze A, Thiagarajah MC, Sathian K. 1999. Tactile perception in developmental dyslexia: a psychophysical study using gratings. Neuropsychologia 37:1201–1211.

Han G, Choi S. 2010. Extended rate-hardness: A measure for perceived hardness. Lect Notes Comput Sci 6191 LNCS:117–124. doi:10.1007/978-3-642-14064-8_18

Hipp J, Arabzadeh E, Zorzin E, Conradt J, Kayser C, Diamond ME, Konig P. 2006. Texture signals in whisker vibrations. J Neurophysiol 95:1792–1799.

Hollins M, Bensmaia SJ, Washburn S. 2001. Vibrotactile adaptation impairs discrimination of fine, but not coarse, textures. Somat Mot Res 18:253–262.

Hollins M, Risner SR. 2000. Evidence for the duplex theory of tactile texture perception. Percept Psychophys 62:695–705.

Hubel DH, Wiesel TN. 1968. Receptive fields and functional architecture of monkey striate cortex. J Physiol 195:215–243.

Hyllienmark L, Brismar T, Ludvigsson J. 1995. Subclinical nerve dysfunction in children and adolescents with IDDM. Diabetologia 38:685–692.

Jadhav SP, Wolfe J, Feldman DE. 2009. Sparse temporal coding of elementary tactile features during active whisker sensation. Nat Neurosci 12:792–800.

Johansson RS, Birznieks I. 2004. First spikes in ensembles of human tactile afferents code complex spatial fingertip events. Nat Neurosci 7:170–7. doi:10.1038/nn1177

Johansson RS, Westling G. 1987. Signals in tactile afferents from the fingers eliciting adaptive motor responses during precision grip. Exp Brain Res Res 66:141–154. doi:10.1007/BF00236210

Jones LM, Depireux DA, Simons DJ, Keller A. 2004. Robust temporal coding in the trigeminal system. Science (80-) 304:1986–1989.

Laasonen M, Service E, Virsu V. 2001. Temporal order and processing acuity of visual, auditory, and tactile perception in developmentally dyslexic young adults. Cogn Affect Behav Neurosci 1:394–410.

LaMotte RH, Mountcastle VB. 1975. Capacities of humans and monkeys to discriminate vibratory stimuli of different frequency and amplitude: a correlation between neural events and psychological measurements. J Neurophysiol 38:539–559. doi:psychophysics; monkeys; haptic; vibration; tactile; vibro-tactile; frequency; amplitude; integrated measure

Lawrence DA, Pao LY, Dougherty AM, Salada MA, Pavlou Y. 2000. Rate-hardness: a new performance metric for haptic interfaces. IEEE Trans Robot Autom 16:357–371. doi:10.1109/70.864228

Luna R, Hernandez A, Brody CD, Romo R. 2005. Neural codes for perceptual discrimination in primary somatosensory cortex. Nat Neurosci 8:1210–1219.

Manfredi LR, Saal HP, Brown KJ, Zielinski MC, Dammann 3rd JF, Polashock VS, Bensmaia SJ. 2014. Natural scenes in tactile texture. J Neurophysiol 111:1792–1802. doi:10.1152/jn.00680.2013

Maravall M, Petersen RS, Fairhall AL, Arabzadeh E, Diamond ME. 2007. Shifts in coding properties and maintenance of information transmission during adaptation in barrel cortex. PLoS Biol 5:e19.

Oladazimi M, Brendel W, Schwarz C. 2018. Biomechanical Texture Coding in Rat Whiskers. Sci Rep 8:11139. doi:10.1038/s41598-018-29225-9

Oldfield RC. 1971. The assessment and analysis of handedness: The Edinburgh inventory. Neuropsychologia 9:97–113. doi:10.1016/0028-3932(71)90067-4

Petersen RS, Brambilla M, Bale MR, Alenda A, Panzeri S, Montemurro MA, Maravall M. 2008. Diverse and temporally precise kinetic feature selectivity in the VPm thalamic nucleus. Neuron 60:890–903.

Pruszynski JA, Johansson RS. 2014. Edge-orientation processing in first-order tactile neurons. Nat Neurosci 17:1404–9. doi:10.1038/nn.3804

Ritt JT, Andermann ML, Moore CI. 2008. Embodied information processing: vibrissa mechanics and texture features shape micromotions in actively sensing rats. Neuron 57:599–613. doi:10.1016/j.neuron.2007.12.024

Salinas E, Hernandez A, Zainos A, Romo R. 2000. Periodicity and firing rate as candidate neural codes for the frequency of vibrotactile stimuli. J Neurosci 20:5503–5515.

Schwarz C. 2016. The slip hypothesis: Tactile perception and its neuronal bases. Trends Neurosci 39:449–462. doi:doi: 10.1016/j.tins.2016.04.008

Skedung L, Arvidsson M, Chung JY, Stafford CM, Berglund B, Rutland MW. 2013. Feeling small: exploring the tactile perception limits. Sci Rep 3:2617. doi:10.1038/srep02617

Tong J, Ngo V, Goldreich D. 2016. Tactile length contraction as Bayesian inference. J Neurophysiol 116:369–379. doi:10.1152/jn.00029.2016

Verrillo RT, Bolanowski SJ, McGlone FP. 1999. Subjective magnitude of tactile roughness. Somatosens Mot Res 16:352–360. doi:10.1080/08990229970401

Waiblinger C, Brugger D, Schwarz C. 2015a. Vibrotactile discrimination in the rat whisker system is based on neuronal coding of instantaneous kinematic cues. Cereb Cortex 25:1093–106. doi:10.1093/cercor/bht305

Waiblinger C, Brugger D, Whitmire CJ, Stanley GB, Schwarz C. 2015b. Support for the slip hypothesis from whisker-related tactile perception of rats in a noisy environment. Front Integr Neurosci 9:53. doi:10.3389/fnint.2015.00053

Weber AI, Saal HP, Lieber JD, Cheng J-W, Manfredi LR, Dammann JF, Bensmaia SJ. 2013. Spatial and temporal codes mediate the tactile perception of natural textures. Proc Natl Acad Sci 110:17107–12. doi:10.1073/pnas.1305509110

Wichmann FA, Hill NJ. 2001. The psychometric function: I. Fitting, sampling, and goodness of fit. Percept Psychophys 63:1293–1313.

Wolfe J, Hill DN, Pahlavan S, Drew PJ, Kleinfeld D, Feldman DE. 2008. Texture coding in the rat whisker system: slip-stick versus differential resonance. PLoS Biol 6:e215.

Yoshioka T, Gibb B, Dorsch AK, Hsiao SS, Johnson KO. 2001. Neural coding mechanisms underlying perceived roughness of finely textured surfaces. J Neurosci 21:6905–6916.

